# Effects of Different Strategies for Exploiting Genomic Selection in Perennial Ryegrass Breeding Programs

**DOI:** 10.1101/2020.05.16.099267

**Authors:** Hadi Esfandyari, Dario Fè, Biructawit Bekele Tessema, Lucas L. Janss, Just Jensen

## Abstract

Genomic selection (GS) is a potential pathway to accelerate genetic gain for perennial ryegrass *(Lolium perenne* L.). The main objectives of the present study were to investigate the level of genetic gain and accuracy by applying GS in commercial perennial ryegrass breeding programs. Different scenarios were compared to a conventional breeding program. Simulated scenarios differed in the method of selection and structure of the breeding program. Two scenarios (Phen-Y12 and Phen) for phenotypic selection and three scenarios (GS-Y12, GS and GS-SP) were considered for genomic breeding schemes. All breeding schemes were simulated for 25 cycles. The amount of genetic gain achieved was different across scenarios. Compared to phenotypic scenarios, GS scenarios resulted in a significantly larger genetic gain for the simulated traits. This was mainly due to more efficient selection of plots and single plants based on GEBV. Also, GS allows for reduction in cycle time, which led to at least a doubling or a trebling of genetic gain compared to the traditional program. Reduction in additive genetic variance levels were higher with GS scenarios than with phenotypic selection. The results demonstrated that implementation of GS in ryegrass breeding is possible and presents an opportunity to make very significant improvements in genetic gains.

## Introduction

Perennial ryegrass *(Lolium perenne* L.) is one of the most cultivated forage species in temperate grasslands, mainly farmed for its re-growth capacity after defoliation, and high value as feed for ruminants, due to palatability, digestibility, and nutritive contents (Wilkins, 1991; Fulkerson et al., 1994; Tallowin et al., 1995). Perennial ryegrass is an obligate allogamous species with genetic gametophytic self-incompatibility, and is bred in genetically heterogeneous families (Cornish et al., 1979).

Recurrent selection is currently the most common strategy employed in ryegrass breeding. Such selection mainly rely on phenotypic records for key traits, combined with pedigree and progeny information (Humphreys, 2005). A breeding cycle may include several selection steps based on information on individual plants and/or plots. Breeding cycles are typically long (10–14 yr.), because phenotypes for many key traits (such as dry matter yield and persistency) can only be reliably measured in plot conditions over multiple years, required to assess the effects of competition among plants (Hayes et al., 2013) and to control for genotype by environment (including year) interactions. The most efficient conventional selection schemes for ryegrass achieve an approximate genetic gain of between 0.5 and 0.7% per year for dry matter yield (Wilkins and Humphreys, 2003).

Genomic selection (GS) is a potential pathway to accelerate genetic gain for perennial ryegrass by reducing the length of the breeding cycle as well as increasing selection accuracy (Meuwissen et al., 2001; Hayes et al., 2013; Lin et al., 2014). One of the advantages of GS is that genetic gain can be increased by decreasing the generation interval, as breeding values can be estimated at an early stage (as soon as DNA can be extracted). Application of GS first requires derivation of a prediction equation using both the phenotypes and genotypes of genome-wide distributed markers (usually based on single-nucleotide polymorphisms [SNPs]) measured in a reference population. Genomic estimated breeding values can then be calculated for selection candidates based on genotypes only, and for phenotyped families genomic information will also enhance accuracy of predicted breeding values over the accuracy obtained from own phenotypic records.

Ryegrass breeding programs typically follow the following steps to develop new varieties: (1) parental individuals, selected from elite varieties, are crossed to generate *F*_1_ progenies, (2) seeds from each *F*_1_ are multiplied in isolation to generate *F*_2_ families that are then phenotyped in several replicates and locations as family pools, (3) single plants (SPs) from selected *F*_2_ families are evaluated as individual genotypes, (4) synthetic varieties (SYNs) are constructed by polycrossing several SPs from the best performing *F*_2_ families (generally between 6 and 10 parents), (5) SYNs are maintained and evaluated as family pools, and after selection, (7) the best-performing SYNs are submitted for official testing (Detailed reviews of breeding methods for grasses are presented by Vogel and Pedersen (2010) and Hayes et al. (2013)). Several studies have developed genomic predictions for some traits of perennial ryegrass using information from different stages of the mentioned breeding program. Fè et al. (2016) explored GS for seed production related traits, forage quality and crown rust resistance in commercial germplasm achieving moderate correlations between average phenotypes and GEBVs in the range of 0.2–0.56. Fè et al. (2015) considered the trait of heading date, and using a cross-validation scheme achieved correlations between average phenotype and GEBVs ranging from 0.52 to 0.9. Grinberg et al. (2016) reported high predictive abilities for water-soluble carbohydrate (0.59), dry matter yield (0.41) based on data from previous generations (containing parental genotypes) to predict the performance of derived half-sib populations using genomic best linear unbiased prediction (GBLUP) and machine learning models. Predictive ability for crown rust resistance on individual plants in a large perennial ryegrass population reached a maximum of 0.52 in a study by Arojju et al. (2018). Although GP has reportedly succeeded in ryegrass, however, application of GS into practical breeding schemes is still under development and careful considerations on steps to be improved by GS are needed.

Different breeding programs with or without GS can be compared by computer simulation before empirical application (Lin et al., 2016, 2017b). Based on genetic principles and parameters informed by empirical data, different breeding strategies can be simulated to predict their performance in term of genetic gain, inbreeding and maintenance of genetic variance. For dairy cattle (Bos Taurus), for which GS has perhaps been most successful, computer simulation was first used to demonstrate the benefits of this technology (Schaeffer, 2006). Thus, the main objective of the present study was to investigate the level of genetic gain and accuracy by applying GS in commercial perennial ryegrass breeding programs. This was achieved by first simulating a conventional ryegrass phenotype-based breeding program and then simulating potential entry points and strategies for GS in the breeding program.

## Materials and Methods

### Simulation outline

The simulation study consisted of the following main steps: (i) simulation of ryegrass base population and initial ryegrass varieties, (ii) simulation of conventional breeding and GS schemes in various scenarios.

### Simulation of the base population and initial varieties

The QMSim software (Sargolzaei and Schenkel, 2009) was used to simulate a historical population of 2000 generations with a constant size of 2000 individuals for 1000 generations, followed by a gradual decrease in population size from 2000 to 1000 to create initial linkage disequilibrium (LD). Random mating with replacement was applied across historical generations. In the next step, to simulate the initial varieties, 20 random samples of 200 individuals were drawn from the last generation of the historical population and, within each sample, individuals were randomly mated for another one generation for variety formation.

### Genome

A genome consisting of 7 chromosome of 100 cM with 100 segregating QTL and 1000 SNPs per chromosome was simulated (Table 1). The bi-allelic genotype at each locus was represented by 0 (homozygous), 1 (heterozygous) or 2 (homozygous alternative allele). Both QTL and SNPs were randomly distributed over the chromosomes. In each meiosis, the number of recombination per chromosome were sampled from a Poisson distribution (λ = 1). To obtain the required number of segregating loci after 2000 generations, about two to three times as many bi-allelic loci were simulated by sampling initial allele frequencies from a uniform distribution and applying a recurrent mutation rate of 2.5 × 10^-5^. Mutation rates of loci were determined on the basis of the number of polymorphic loci in generation 2000 of the preliminary analysis that were necessary to obtain 1000 polymorphic SNPs and 100 QTL per chromosome. Mutations were limited to the loci in historical population. SNPs and QTL were distinct loci and were randomly drawn from segregating loci, with a minor allele frequency (MAF) higher than 0.05, in generation 2000.

**TABLE 1.**
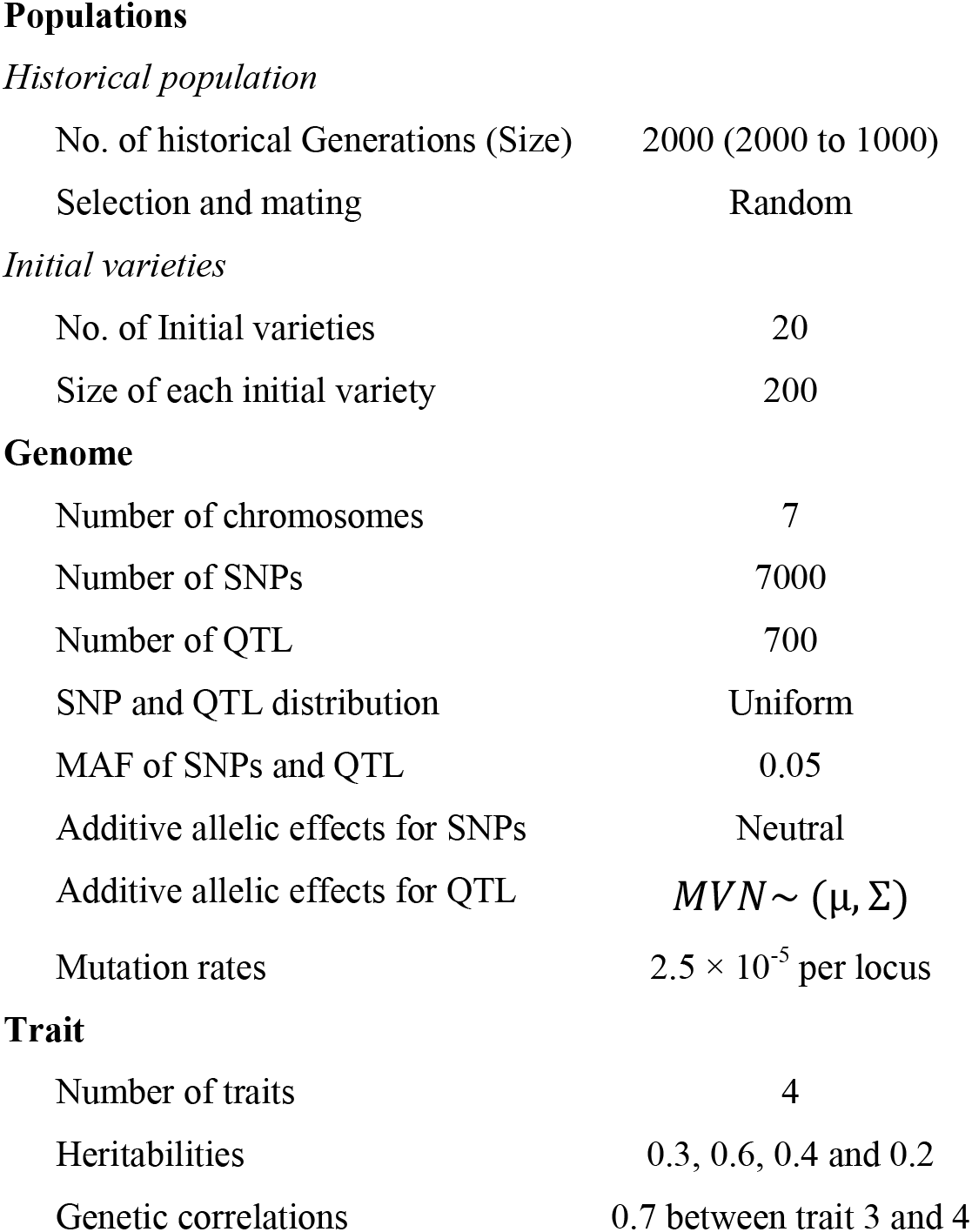
Parameters of the simulation process

### Simulation of true breeding values and phenotypes for traits

Four traits were simulated: Trait 1 (*h*^2^ = 0.3), Trait 2 (*h*^2^ = 0.6), Trait 3 (*h*^2^ = 0.4), and Trait 4 (h^2^ = 0.2). Arbitrary traits with different heritabilities were considered to reflect the heritability estimates by empirical studies for some economically important traits (e.g., forage yield and rust resistance) in ryegrass breeding (Fé et al., 2015, Wang et al., 2011). For all traits, the plot heritability was considered which theoretically is equal to the square of the correlation between the sum (mean) of TBV of individuals and sum (mean) of the phenotypes of individuals in the plots. So, by trial and error additive variance (additive effects *per se*) was calibrated in a way that the realized *h*^2^ plot was equal to the desired *h*^2^ plot.

True breeding values for the traits were generated as follows. Allele substitution effects for quantitative trait loci (QTL) were sampled from a multivariate normal distribution *MVN~* (*μ*,∑), where *μ* = 0 for all traits and Σ is a covariance matrix among traits (see below). Each trait had 700 QTL randomly drawn from segregating loci across the whole genome with MAF > 0.05. All 700 QTL were shared (pleiotropic) across all traits. The QTL effects were sampled with a covariance of ~0.2 for Trait 3 and Trait 4 and 0 for all other pairs of traits. As a result, genetic correlation between Trait 3 and Trait 4 was 0.7, while there was no genetic correlation among other traits. The TBVs for each trait were calculated as follows:

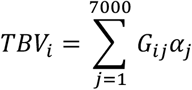

where *G_ij_* is the genotype (taking values of 0, 1, and 2) of individual *i* at locus *j*, and *α_j_* is the QTL allele substitution effect. The TBV of a plot was approximated as the average TBV across all individuals in the plot.

All QTL effects were assumed to be additive and phenotypes were generated by adding a random normal deviate 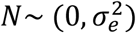 to TBVs where 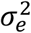 for each trait was equal to deviation of additive variance from the phenotypic variance of each trait (i.e., 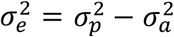).

### One cycle of conventional breeding program

All stages of a typical 12-yr conventional breeding scheme are shown in Figure 1. Breeding scheme commences with 20 initial varieties from the base population. Two randomly selected plants from a pair of varieties were randomly mated to create an *F*_1_ family of full sibs. The procedure was repeated to create 250 *F*_1_ families of 40 plants in each family (10,000 individuals in total), thus, on average 25 plants were chosen as parents from each of the initial varieties. Plants within each *F*_1_ family were randomly mated to produce both the *F*_2_ families that were grown in plots and *F*_2_ single plants in greenhouse (by a delay of one year). This means for each *F*_2_ family single plants were also available for later use (see below). Plants in each *F*_2_ family were the result of random mating among full sib plants within each *F*_1_ family with absence of selfpollination (ensured by self-incompatibility). In the next step, 50 *F*_2_ families with high performance in the three trait phenotypes (Tr. 1, Tr. 3 and Tr. 4) were identified (Equation 1) and their corresponding single plants in green house were used for creation of synthetic (SYN) groups as following. From the 40 single plants belonging to each top 50 *F*_2_ family, 8 single plants were randomly chosen to be used as parents of SYN groups (400 single plants in total). Selected single plants were grouped into eight-parent synthetics (50 groups) by their similarity of the heading time phenotypes that was Trait 2 (*h*^2^ = 0.6) in our simulation. Three categories were considered for heading time phenotypes as early, intermediate and late. In each category, there were ~50/3 eight-parent groups on average. In the next step, the 8-parent groups were polycrossed (SYN1), followed by random mating within each synthetic group. This step in practice is mainly performed to obtain sufficient seed for SYN2 plot establishment. In the final step, 20 SYN2 plots were selected using a selection index (Equation 1) based on performance in Tr. 1, Tr. 3 and Tr. 4, followed by random mating within each SYN2 to created SYN3 plots.

**FIGURE 1.**
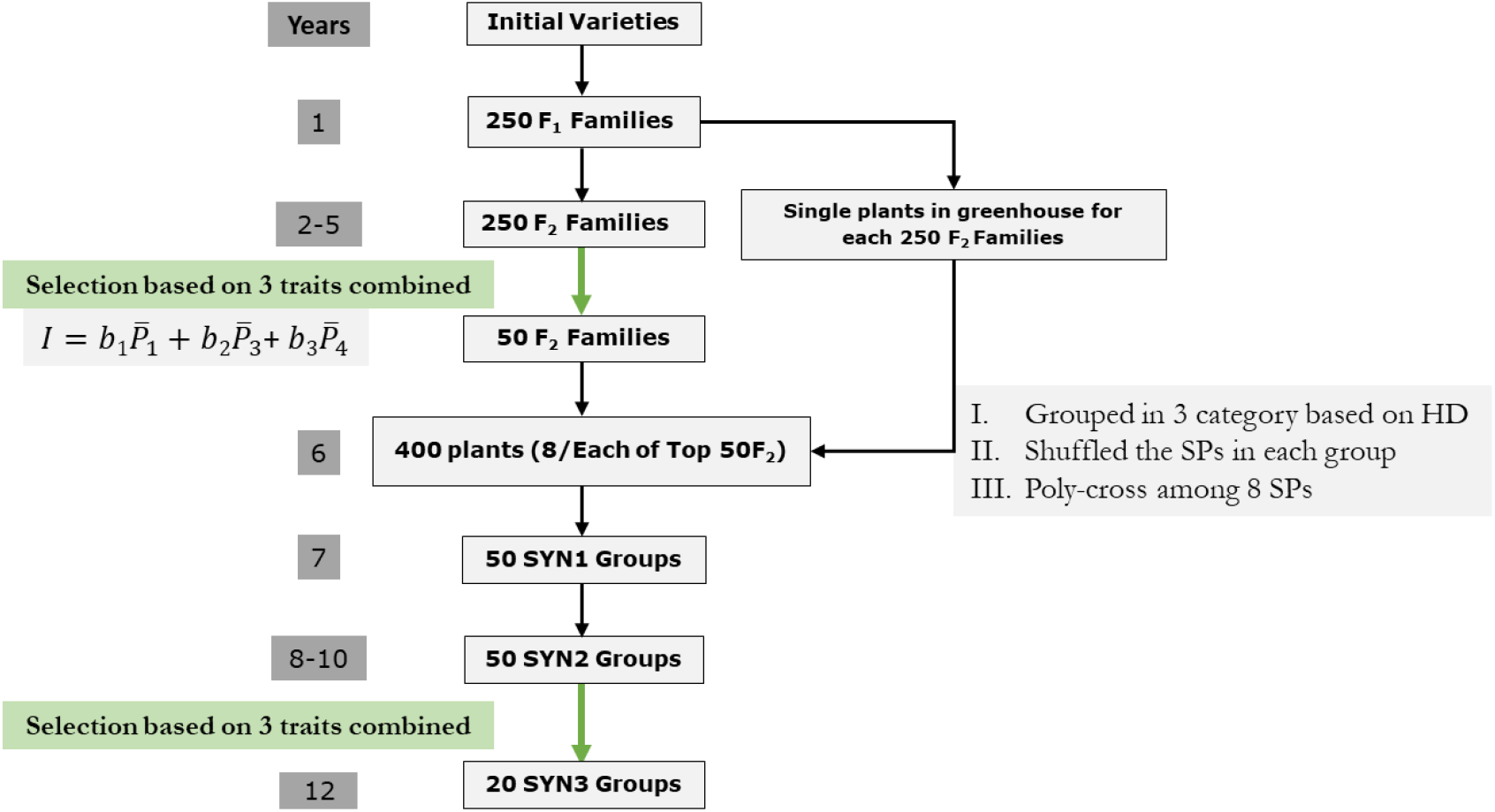
Schematic of one cycle in the conventional (phenotypic) breeding strategy of perennial ryegrass. Green arrows indicate selection stages. HD, heading date; SP, single plants.

The conventional breeding program simulated in the present study included two rounds of multi-trait selection. First round of selection was at the stage of *F*_2_ families and the second round of selection was in SYN2. Multi-trait selection considered Tr. 1, Tr. 3 and Tr. 4 simultaneously using a selection index as follows:

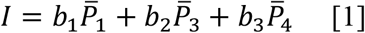

where *b*_1_, *b*_2_ and *b*_3_ are the selection index weights for Tr. 1, Tr. 3 and Tr. 4 and 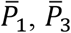 and 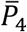 are the mean phenotypes of the three traits in plots, respectively. All *b_i_*, were set to 1/3 to achieve standardized emphasis on each trait. It should be noted that this selection index does not reflect the multi-trait selection in practice as that would include defining economic weights and computing *b*-values based on genetic and phenotypic variances and covariances. Nevertheless, the emphasis on Trait 3 was still slightly greater as a result of the higher *h*^2^ of this trait. Trait 2 was assumed as heading date and was only used for grouping single plants into groups with similar heading date, thus single plants were not selected for this trait.

### One cycle of genomic selection breeding program

The proposed GS-based breeding schemes was designed to integrate with the current breeding program by replacing phenotypic selection points with GS (Figure 2). All steps of the genomic breeding program were similar to the conventional breeding with some modifications as follows. Similar to phenotypic selection, each cycle started by crossing of single plants from initial varieties to create 250 *F*_1_ families and random mating among full-sibs within each *F*_1_ family, which resulted in 250 *F*_2_ families. At this stage, compared to phenotypic breeding program, single plants in green house were not planted for all *F*_2_ families. Instead, once top 50 *F*_2_ families were selected using a selection index based on GEBV of Tr. 1, Tr. 3 and Tr. 4 (Equation 2), *F*_2_ single plants were planted only for those top 50 *F*_2_ families. In the next step, 400 single plants were selected using a selection index (Equation 2) based on combination of GEBV of the three traits. Similar to phenotypic breeding program, selected single plants were grouped into eight-parent groups (50 groups) by their similarity of the heading date phenotypes (Tr. 2). In the next step, the 8-parent groups were polycrossed (SYN1), followed by random mating within each synthetic to create SYN2 groups. In the final step, 20 SYN2 plots were selected using a selection index (Equation 2) based on GEBV of Tr. 1, Tr. 3 and Tr. 4, followed by random mating within each SYN2 to create SYN3 groups. Similar to phenotypic breeding scheme, the GS scheme had two rounds of multi-trait selection. First round of selection was at the stage of *F*_2_ families and the second round of selection was in SYN2 plots step. The selection of the top plots in both stages, was based on the following multi-trait index:

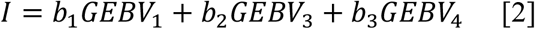

Where *b*_1_, *b*_2_ and *b*_3_ are the selection index weights for Tr. 1, Tr. 3 and Tr. 4 and *GEBV_i_* (i=1, 3 and 4) is the GEBV of the trait in plots. The *b_i_* values were similar to the phenotypic breeding.

Genomic breeding strategy had two major differences with phenotypic selection. First, *F*_2_ single plants were planted only for the selected 50 *F*_2_ families. Second, selection of the 400 *F*_2_ single plants for poly-crossing was not random as phenotypic strategy but based on their GEBV using an index. To do so, for all *F*_2_ single plants, GEBV for the three traits were estimated and the top 400 single plants were selected based on a similar index for the selection of top plots (Equation 2). However, *GEBV_i_* in the Equation 2 was estimated for a single plant rather than based on a plot.

A key aspect of a GS is the establishment of a reference population, which is used to train prediction equations for traits in the breeding goal. The reference population for the estimation of marker effects and calculation of GEBV for *F*_2_ families and *F*_2_ single plants was recruited from *F*_2_ families. For each *F*_2_ family one phenotype and one summary genotype (allele dosage per marker) per plot was generated (similar to Ashraf et al., 2014). In other words, every *F*_2_ family was treated as a proxy individual in the reference population with one phenotype and genotypes of mean dosage of 20 plants. Genotypes for the plots were represented by the allele dosage, which is the mean genotype of a subset of 20 individuals per plot and per SNP. Therefore, genotypes were real numbers between 0 and 2, which calculated from allele frequencies, rather than integers 0, 1, or 2. For example if the alleles at a SNP were A and T, with the T designated as the second allele, and the frequency of the T allele was 0.7 in the plot, the allele dosage would be 0.7*2 = 1.4. Allele dosage or summary genotype was used because each plot contains a large number of individuals each with a different genetic makeup. Once the reference population was established and SNP effects were estimated (Equation 3), GEBV of the 3 traits for each of the selection candidates (*F*_2_ families) was calculated. Here, genotypes of each plot were again represented by mean allele dosage of 20 plants per plot. However, for *F*_2_ single plants, the observed genotype of each single plant was used to estimate GEBV.

**FIGURE 2.**
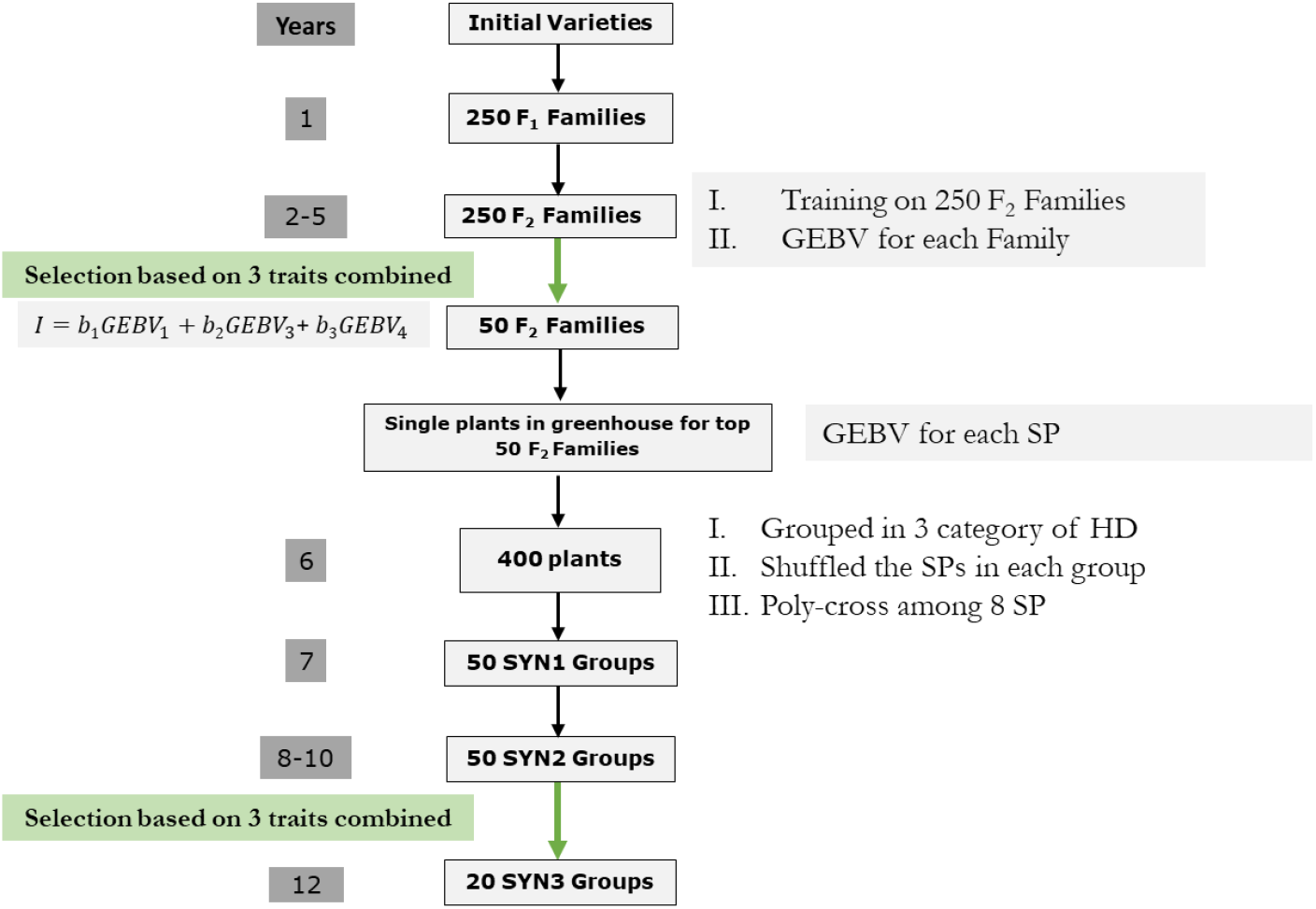
Schematic of one cycle in the genomic selection breeding strategy. Green arrows indicate selection stages. HD, heading date; SP, single plants; GEBV, genomic estimated breeding value.

### Logical flow of the breeding cycles

Both the conventional and genomic breeding schemes were simulated for 36 years by starting a new cycle every year and 36 years of breeding program was equivalent to 25 cycles (Figure 3). Each year, one breeding cycle was started by crossing single plants from available parents to create *F*_1_ families and so on. Each breeding cycle takes 12 years to complete, from *F*_1_ to SYN3 groups.

**FIGURE 3.**
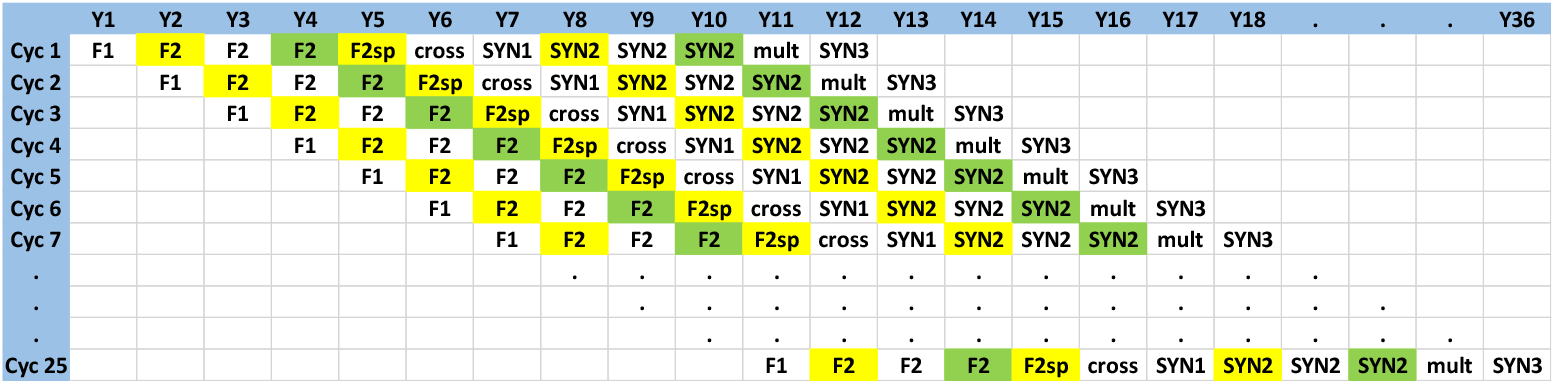
Logical flow of the simulated breeding cycles. Y= year.

### Scenarios

We examined five different scenarios (Table 2). Two scenarios (Phen-Y12 and Phen) were considered for phenotypic selection and three scenarios (GS-Y12, GS and GS-SP) were simulated for genomic breeding schemes. The simulations were performed with scripts developed in R version 3.4.0 (R Core Development Team, 2016). All scenarios were conducted with 50 independent replicates except for the historical population, which was same for all replicates and scenarios.

**TABLE 2.** The five different simulated scenarios.

| Scenarios | Independent cycles<sup>a</sup> | Input from previous cycles |
| --- | --- | --- |
| Phen-Y12 | 11 | SYN2 groups |
| Phen | 5 | $F_2$ single plants, SYN2 groups |
| GS-Y12 | 11 | SYN2 groups |
| GS | 5 | $F_2$ single plants, SYN2 groups |
| GS-SP | 5 | $F_2$ single plants, SYN2 single plants |

<sup>a</sup> Number of cycles where initial varieties were used as parents.

### Sc. Phen-Y12

In this scenario, all cycles for the first 11 years were started by sampling single plants from 20 initial varieties. For each of these cycles, single plants sampled from initial varieties were crossed to create *F*_1_ families, following the same steps mentioned above for conventional breeding program (Figure 1). As new cycle was starting in each year, it was assumed that there was no material exchange among the 11 initial cycles. For the following cycles after cycle 11, the initial material (parents) for making *F*_1_ families were recruited from output (SYN2) of previous cycles. As an example, for the start of cycle 12, the SYN2 groups of cycle 1 were already available and served as a starting material for cycle 12. Similarly, cycle 13 could start with SYN2 groups of cycle 1 and 2, which were available at the starting time of cycle 13 (Figure 3). Finally, for cycle 25, SYN2 groups of cycle 1 to 14 were available to be used as initial materials. This reflected the outcomes of a breeding practice in which elite varieties serve as the starting material for multiple breeding cycles. As each cycle required 12 years from the stage of *F*_1_ families to SYN3 groups to be accomplished, 36 years of breeding program was equivalent to 25 full cycle of breeding and selection in this scenario.

### Sc. Phen

In this scenario, the 5 initial cycles were similar to the 11 initial cycles of Sc. Phen-Y12 meaning that a new cycle started by sampling and crossing single plants from the base population. However, for this scenario, it was assumed that both *F*_2_ single plants and SYN2 groups could be used as parents once they are available. So, for starting cycle 6 as *F*_2_ single plants from cycle 1 were already available, they were used as the starting material for cycle 6. Similarly, cycle 7 could start with crossing of *F*_2_ single plants from cycle 1 and 2. In other words, once *F*_2_ single plants from earlier cycles were available, they served as initial material for the later cycles. For breeding cycle 12, besides *F*_2_ single plants from previous cycles SYN2 groups of cycle 1 could also be used for making crosses. In this scenario, all available *F*_2_ single plants and SYN2 groups from previous cycles could be used as starting material for a new cycle. Compared to Sc. Phen-Y12, in which 11 years were needed to start a new cycle by the output of cycle 1, in Sc. Phen using *F*_2_ single plants as starting material upon their availability, could potentially reduce the breeding cycle from 12 years to 6 years with the remaining 6 years primarily used for produce development before marketing.

### Sc. GS-Y12

The breeding structure of this scenario was similar to Sc. Phen-Y12 and can be considered as genomic variant of Sc. Phen-Y12. Similar to Sc. Phen-Y12, the first 11 cycles started by crossing of initial varieties as parents. Once SYN2 groups of cycle 1 were available, they were used as parents for starting cycle 12 and for starting cycle 13, SYN2 groups of both cycle 1 and 2 were available to be used as parents. Thus, in this scenario, only SYN2 groups could be used as starting material for a new cycle once they were available (cycle 12 onward). Similar to Sc. Phen-Y12, first 11 breeding cycles were running independently with no material exchange among the cycles. However, as selection at the stage of *F*_2_ families and *F*_2_ single plants in each cycle were based on GEBV, reference population for estimating marker effects were not limited to the *F*_2_ families of the corresponding cycle. Instead, as breeding program was running over the years, the reference population (*F*_2_ families) were increased by adding new phenotypes and genotypes from the *F*_2_ families across years. This means that for cycle 1 only *F*_2_ families of cycle 1 were used as a reference, however, for cycle 2, *F*_2_ families from both cycle 1 and 2 were used for estimation of marker effects, i.e. every year the size of reference population was growing by 250. Also, as in genomic breeding schemes, selection of SYN2 groups in each cycle were based on GEBV, the reference population for the estimation of marker effects at this stage was not limited to the SYN2 groups of the corresponding cycle. Here, once SYN2 groups were available, phenotype and genotype of each SYN2 group were also added to the same reference population that was used for the estimation of marker effects to be used in calculation of GEBV in *F*_2_ families. In other words, the GEBV for SYN2 groups of cycle 1 were calculated based on a reference population consisting of *F*_2_ families from cycle 1 to 8 and SYN2 groups of cycle 1 and 2. It should be noted that it was assumed SYN2 groups of cycle 1 were selected after being sown in multiplication plots in year 11 and by that time SYN2 groups of cycle 2 could be genotyped to be added to the reference population.

### Sc. GS

This scenario is genomic variant of Sc. Phen. Similar to Sc. Phen, the first 5 cycles started by using initial varieties as parents. For starting cycle 6, *F*_2_ single plants from cycle 1 were available and were used as input material (parents) for this cycle. Similarly, cycle 7 could use *F*_2_ single plants from cycle 1 and 2 as parents for making *F*_1_ families. In this scenario, it was assumed that both *F*_2_ single plants and SYN2 groups could be used as parents of a new cycle once they were available. All steps within each cycle of this scenario was also similar to Sc. GS-Y12. However, compared to Sc. GS-Y12, where only SYN2 groups could be used as starting material after 11 years, in this scenario using *F*_2_ single plants of previous cycles, upon their availability, resulted in reduction of breeding cycle. Construction of the reference population were similar to Sc. GS-Y12. As breeding program was running, genotypes and phenotypes from *F*_2_ families and SYN2 groups were added to the reference population to estimate marker effects. The estimated effects then could be used for calculation of GEBV of the relevant stage.

### Sc. GS-SP

This scenario was the same as Sc. GS with one modification at the stage of SYN2. For this scenario, it was assumed that not only *F*_2_ families have single plants planted in greenhouse, but also SYN2 groups in each cycle were assumed to have SYN2 single plants planted in greenhouse. So, rather than using *F*_2_ single plants and SYN2 groups as parents for making new cycles as it was done in Sc. GS, in Sc. GS-SP, both *F*_2_ single plants and SYN2 single plants could be used as parents for making *F*_1_ families for new cycles. Similar to *F*_2_ single plants, GEBV for SYN2 single plants was estimated based on observed genotypes of single plants rather than mean allele dosage of SYN2 groups. The construction of reference population in this scenario was similar to GS-Y12 and GS scenarios, where *F*_2_ families and SYN2 groups were added to the reference population once they become available, resulting in a step wise increase in the size of reference population for the later cycles of breeding program.

## Statistical Analysis

### Estimation of marker effects

Bayesian ridge regression (BRR) implemented in the BGLR package was used to predict effects of SNPs (Perez and de los Campos, 2014). The following model was used to predict the additive effects associated with each SNP:

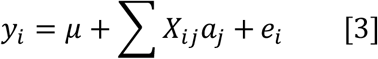

where *y_i_* is the phenotypic observation of plot *i* (*F*_2_ family or SYN2 group) in the training data, *μ* is the overall mean, *X_ij_* is the mean allele dosage of 20 plants randomly sampled per plot for marker *j*, ranged from 0 to 2, *a,j* is the random unknown allele substitution effect for marker *j,* and *e_t_* is the residual effect for plot *i,* and Σ denotes summation over all SNPs *j*. For BRR model Gaussian priors for the marker effects are assumed. The BGLR package assigns scaled-inverse *χ*^2^ densities to the variance parameters whose hyperparameters were given values using the default rules implemented in BGLR, which assign 5 degrees of freedom and calculates the scale parameter based on the sample variance of the phenotypes (further details can be found in Perez and de los Campos, (2014)). For each analysis, the Gibbs sampler was run for 50,000 iterations, with the first 10000 discarded as burn in.

Selections of single plants in the genomic breeding schemes at the stage of *F*_2_ or SYN2 were based on the GEBVs. Thus, from the estimates of allele substitution effects *(â)*, GEBV for each trait in all single plants was calculated as:

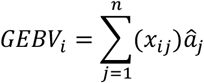

Where *x_ij_* is the copy number of a given allele of marker *j*, coded 0, 1 and 2 for *aa, aA* and *AA*, respectively and Σ denotes summation over all SNPs *j*. GEBV for *F*_2_ families or SYN2 groups was calculated similarly except that *X_ij_* represented the mean allele dosage of 20 plants randomly sampled per plot for marker *j*.

### Genetic gain

Genetic gain in all scenarios was investigated by the cumulative genetic standard deviations (Δ_*G*_) in later cycles after cycle 1, which were calculated as follows:

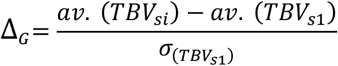

where *av*. (*TBV_si_*) and *av*. (*TBV*_*S*1_) are the mean TBVs of the *F*_1_ groups in cycle 1 and in later cycles (*i* = 2 to 25), and *σ* (*TBV*_*s*1_) is the standard deviation of the *F*_1_ true breeding values at cycle 1.

### Accuracy

Selection accuracy in the genomic breeding schemes for *F*_2_ and SYN2 single plants was evaluated as the Pearson correlation between individual GEBVs and TBVs in each cycle. Selection accuracy at the stage of SYN2 for phenotypic scenarios was calculated as the Pearson correlation between plot phenotypic performance and TBVs, whereas for genomic scenarios, it was the Pearson correlation between plot GEBV and TBV in each cycle.

### Additive genetic variance

Additive genetic variance in all scenarios was measured as the variance of mean TBV of *F*_1_ groups in each cycle.

### Linkage disequilibrium

To evaluate the extent and magnitude of LD in the initial varieties, LD was measured by *r*^2^ and was compared with expected LD with mutation (Tenesa et al., 2007), and to empirically observed LD in ryegrass. Only markers with a MAF greater than 0.05 were considered, because the power of detection of LD between two loci is minimal when at least one of the loci has an extreme allele frequency (Goddard et al., 2000). To determine the decay of LD with increasing distance between SNPs, the average *r*^2^ within each variety was expressed as a function of distance between SNPs. SNP pairs were grouped by their pairwise distance into intervals of 1 cM, starting from 0 up to 20 cM. The average *r*^2^ for SNP pairs in each interval was estimated as the mean of all *r*^2^ within that interval.

## Results

### Genetic gain

The amount of genetic gain achieved was different across scenarios (Figure. 4). Compared to phenotypic scenarios, GS scenarios achieved significantly higher genetic gain (in genetic standard deviation units) for all traits. In all scenarios, as long as cycles were running independently and using initial varieties (base population) as parents, no cumulated genetic gain was observed. However, once the output of previous cycles were available and used as the starting material, a trend in genetic gain could be realized with different rates across scenarios. It should be noted that the comparison of genetic gain among scenarios should take into account the structure of the breeding program, meaning that comparison for genetic gain was made between breeding structures with similar set up. In this sense, Sc. Phen-Y12 and GS-12 can be compared to each other as they have similar breeding structure (starting with 11 independent cycles). Likewise, Sc. Phen, GS and GS-SP can be compared to each other for having 5 initial cycles starting independently from the base population. Sc. GS-Y12 compared to Sc. Phen-Y12 resulted in higher genetic gain. Similarly, Sc. GS and GS-SP were also superior to Sc. Phen. The genetic gain achieved by Sc. GS and GS-SP were similar for all traits.

**FIGURE 4.**
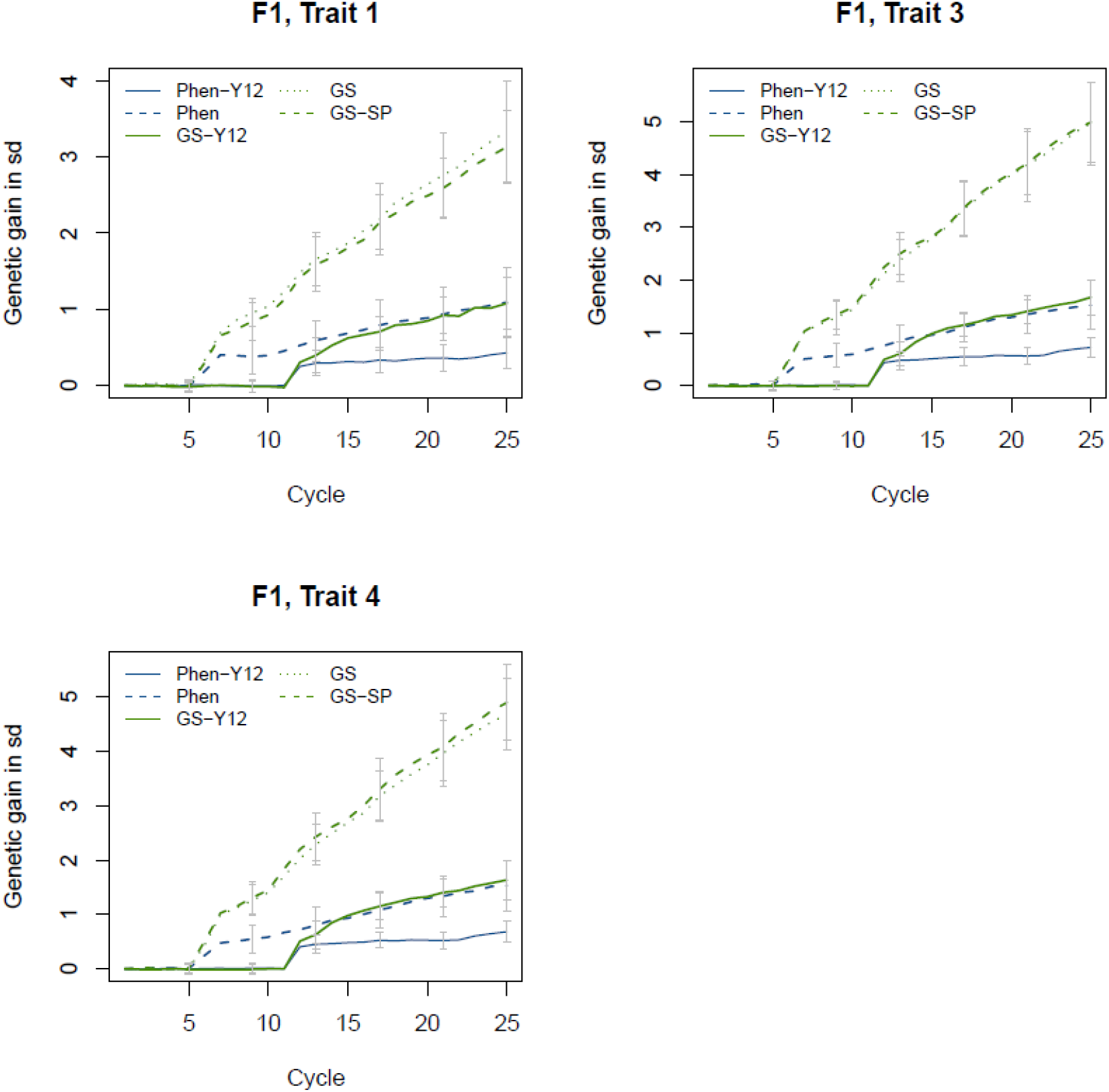
Genetic gain in cumulative genetic standard deviations (sd) for the three traits at the stage of *F*_1_. Error bars indicate standard deviations.

The difference between mean TBV of plots at the stage of and SYN2 within each cycle, for all scenarios, are shown in Figure 5. In all scenarios, mean TBV of SYN2 groups was higher than the mean TBV of *F*_1_ families. The difference between genetic levels of two stages is due to selection (two-step selection) within each cycle. In all scenarios, genetic level of *F*_1_. and SYN2 stages in GS breeding schemes was higher than phenotypic scenarios (Figure 5a). Also, with GS breeding schemes rate of genetic gain within cycle was more than phenotypic scenarios (Figure 5b).

**FIGURE 5.**
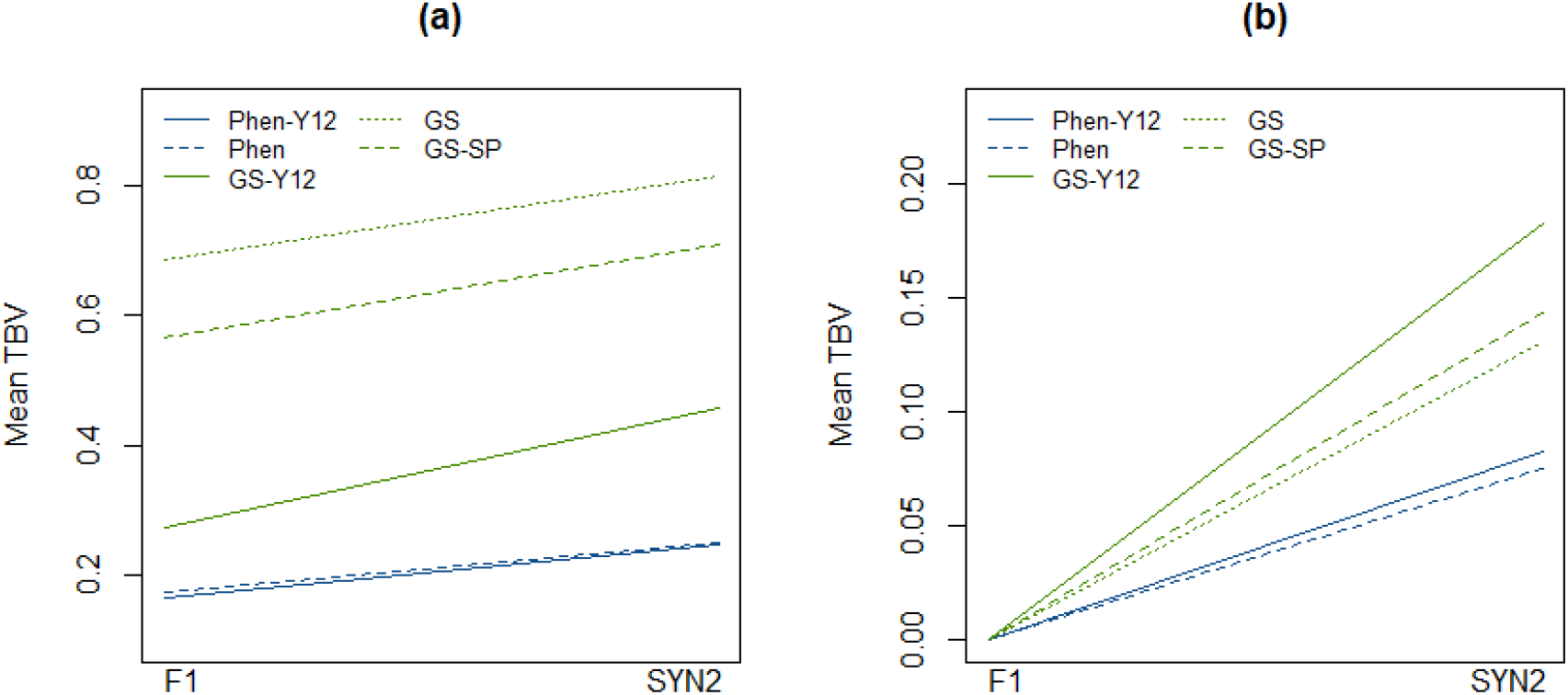
Mean TBV of *F*_1_ and SYN2 stages based on average of 25 cycles. Plot (a) is based on non-scaled and observed values in both stages. In plot (b), mean TBV at stage of *F*_1_ in all scenarios are scaled to zero to illustrate trend of genetic gain within cycles.

### Accuracy

The accuracy of phenotypic and genomic selection for all scenarios for SYN2 is shown in Figure 6. Higher accuracy of selection was obtained for GS scenarios compared to phenotypic scenarios. As expected, for phenotypic scenarios limited accuracies were achieved which were constant across the cycles. However, for GS scenarios accuracy improved across the cycles as a result of increase in the size of the reference population used for the estimation of marker effects. At the end of the breeding program, the highest accuracy was ~0.6 for Sc. GS and GS-SP while the lowest was 0.15 for both phenotype based selection (Phen-Y12 and Phen).

**FIGURE 6.**
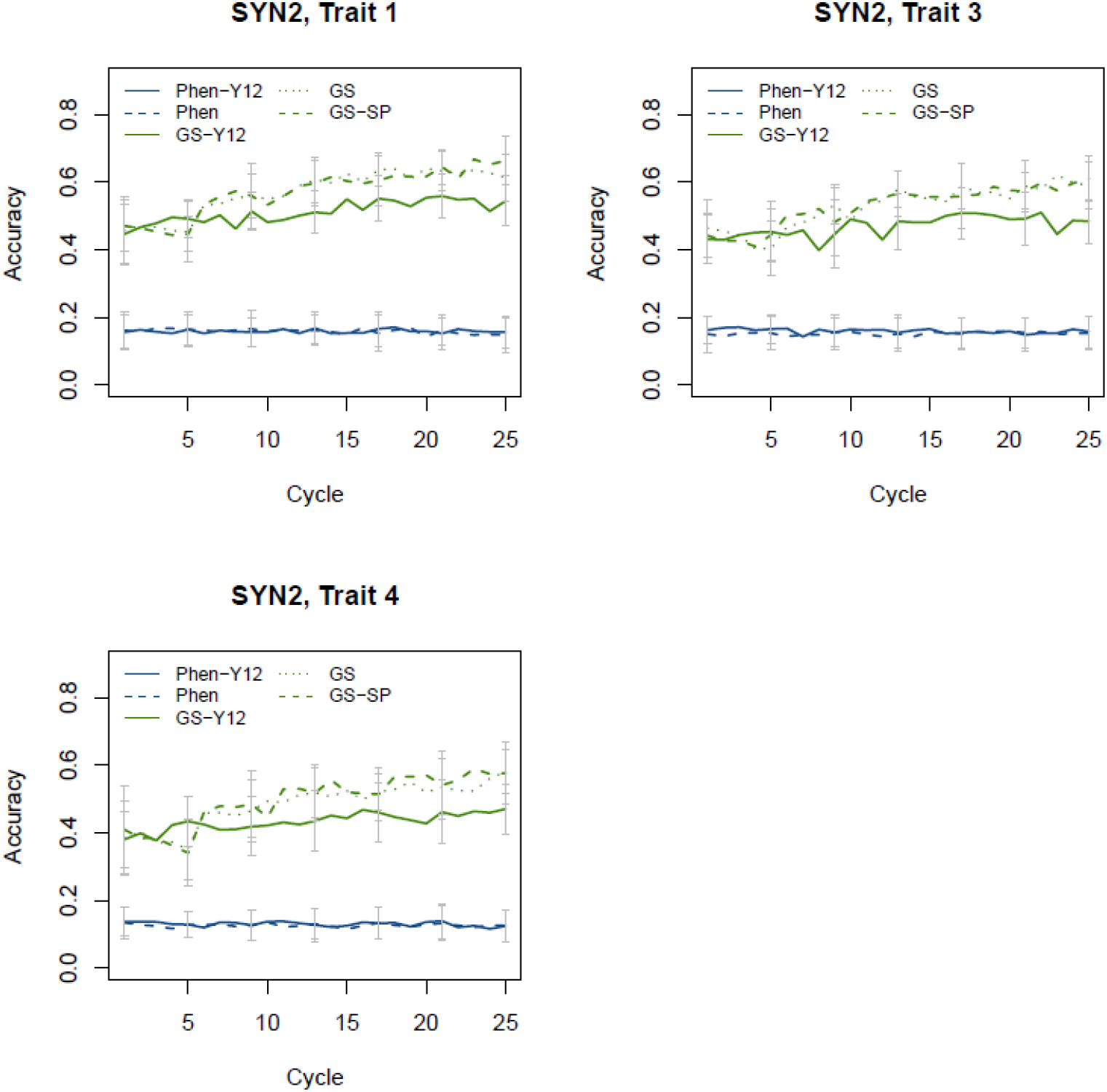
Selection accuracy in SYN2 groups. For genomic scenarios, accuracy was the correlation of GEBV and TBV, while in phenotypic scenario accuracy was the correlation of mean phenotype and TBV. Error bars indicate standard deviations.

The accuracy of GEBVs was evaluated for the three traits at the stage of *F*_2_ and SYN2 single plants in Sc. GS-SP (Figure 7). In both *F*_2_ and SYN2 single plants, accuracies were low (0.15-0.25) for the initial cycles and improved across the cycles as the reference population size increased. For *F*_2_ single plants, Trait 4, had the lowest accuracy as a result of lower *h*^2^. Single plants at the stage of *F*_2_ had higher accuracy that single plants of SYN2. For *F*_2_ single plants, the average accuracy of the three traits for the first and last cycle was 0.25 and 0.55, respectively. The corresponding values for SYN2 single plants were 0.15 and 0.35.

**FIGURE 7.**
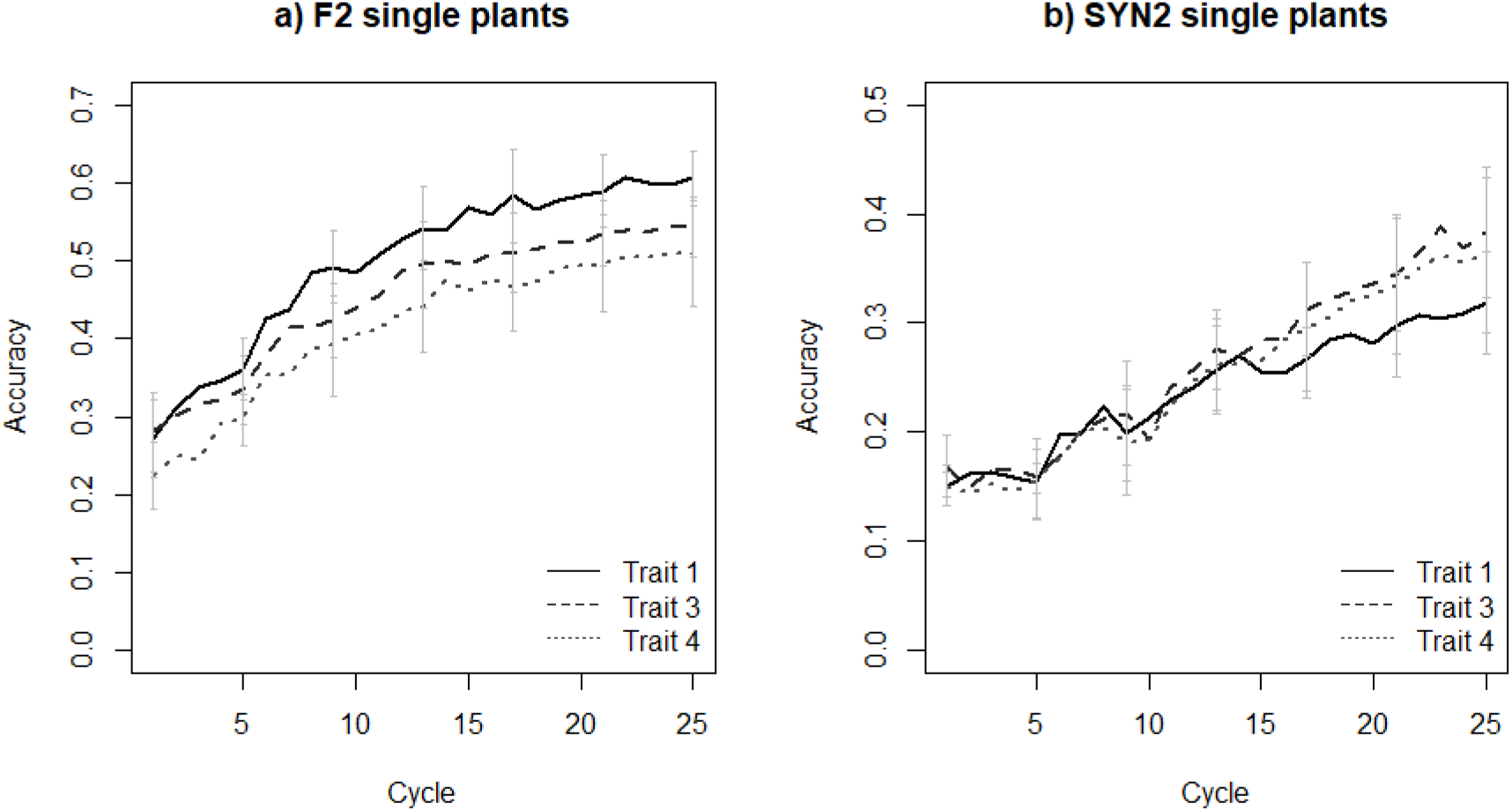
Accuracy of selection in *F*_2_ single plants (plot a) and SYN2 single plants (plot b). Accuracy is the correlation of GEBV and TBV of single plants. Error bars indicate standard deviations.

### Additive genetic variance

Additive genetic variance was measured at the stage of *F*_1_. for all scenarios (Figure 8). Changes in the amount of additive genetic variance was different across scenarios. For phenotypic scenarios, the additive genetic variance was almost constant across the cycles. In GS scenarios, however, additive genetic variance was constant only for the initial cycles (the first 5 cycles for Sc. GS and GS-SP) and was reduced afterwards. At cycle 25, these scenarios retained approximately 60% of the initial additive variance. For Sc. GS-Y12, reduction in the additive variance was less than other genomic scenarios. In fact, for this scenario, the additive variance was constant for the 11 initial cycles as theses cycles were running independently but reduction for variance could be realized for cycle 12 onward.

**FIGURE 8.**
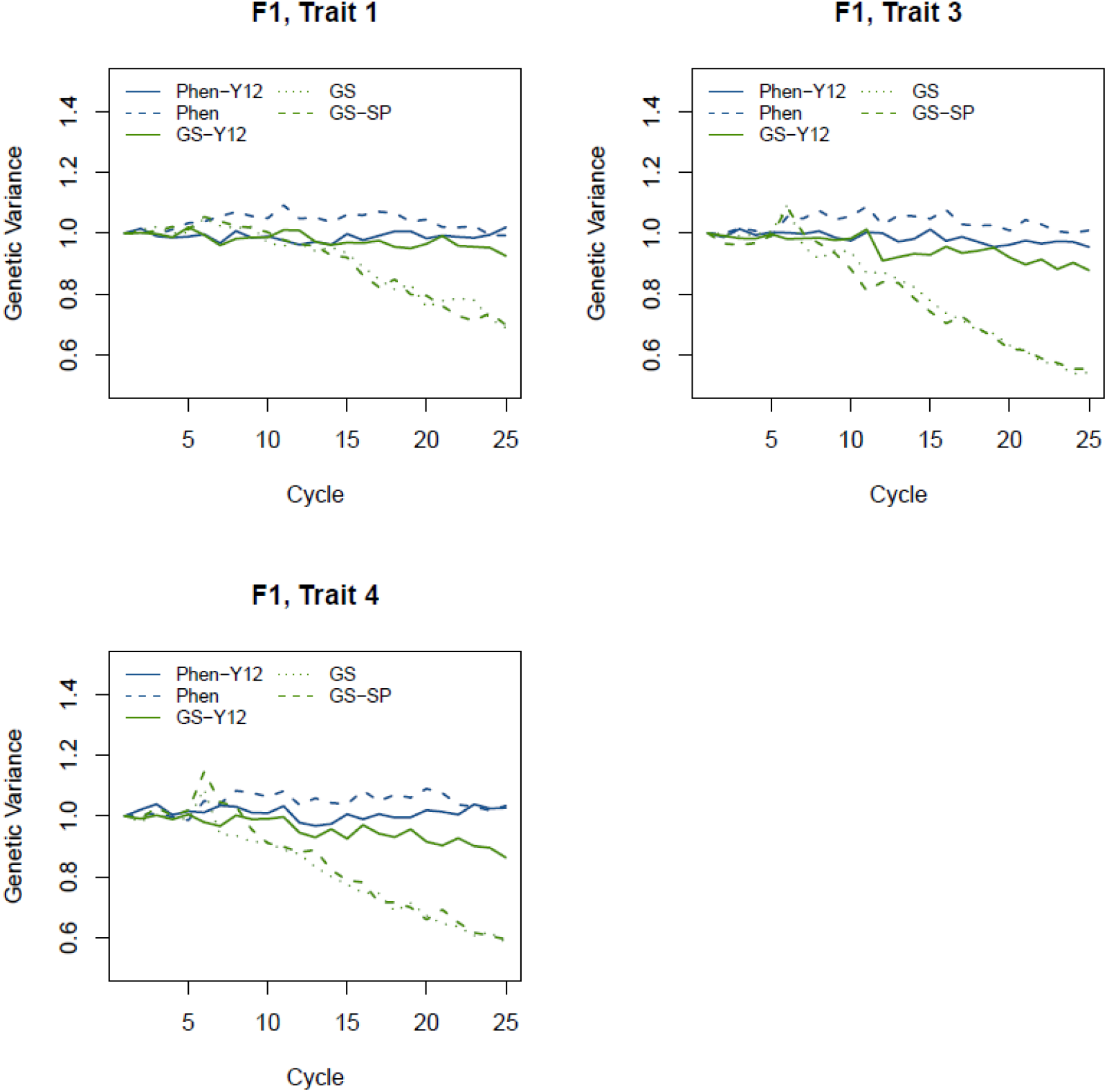
Changes in additive genetic variance in *F*_1_ over the period of 25 breeding cycles. Additive genetic variances for each scenario were standardized by the initial additive variance (cycle 1) of each scenario. Standard deviations ranged from 0.01 to 0.02.

### Linkage disequilibrium

To estimate LD, we used SNP genotypes in all initial varieties. An average *r*^2^ of 0.21 for adjacent SNPs was found which was similar to the empirical levels of LD reported in the literature (Ponting et al., 2007; Brazauskas et al., 2011). Figure 9 displays an overview of the decline of *r*^2^ over distance. As expected, the most tightly linked SNP pairs had the highest average *r*^2^, and the observed average *r*^2^ decreased rapidly as the map distance increased. The observed heterozygosity in initial varieties (He = 0.34) was slightly below those presented by Brazauskas et al. (2011) (0.40) among ryegrass subpopulations. These observations suggest that the structures of the simulated genomes were similar to those of actual ryegrass genomes.

**FIGURE 9.**
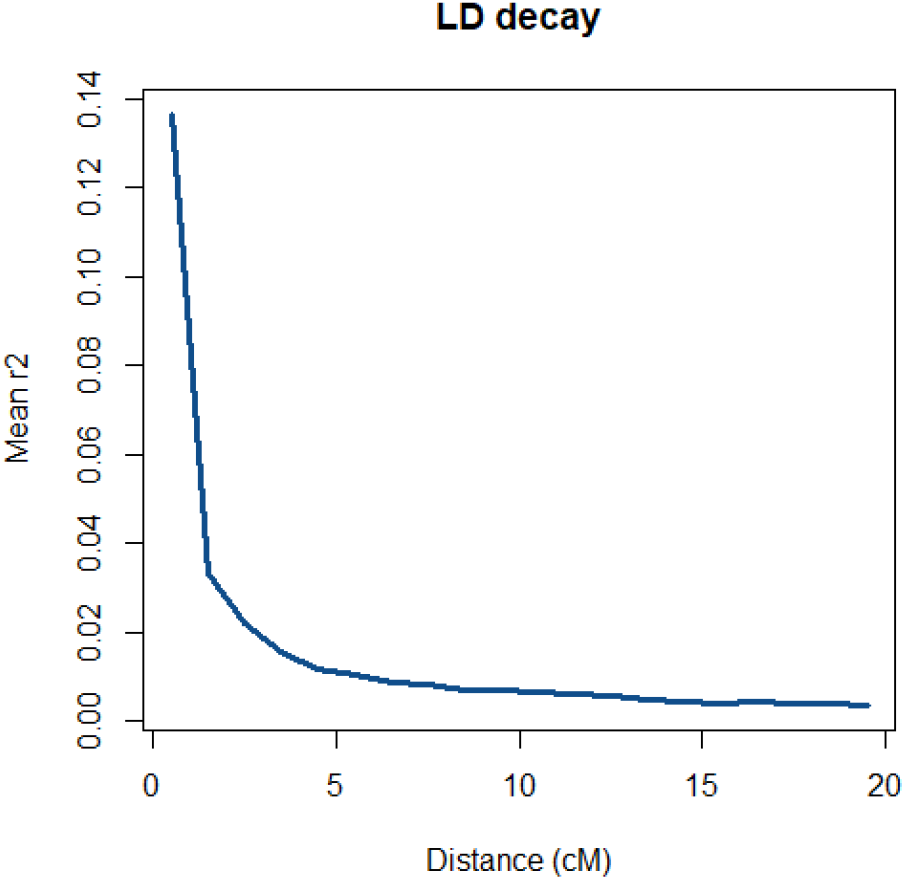
Decay of average *r*^2^ over distance.

## Discussion

The potential effects of applying GS on genetic gain in a commercial ryegrass-breeding program were investigated using simulations. GS breeding schemes resulted in a significantly larger genetic gain for the simulated traits when compared with phenotypic selection. This was mainly due to (a) GS more effectively selecting *F*_2_ single plants as well as *F*_2_ and SYN2 groups using GEBVs (b) GS allowing for reduction in cycle time, which led to at least doubling and trebling genetic gain compared with traditional selection. However, reduction in additive genetic variance levels were higher with GS than with phenotypic selection.

### Genetic Gain

Different scenarios were compared to the conventional breeding program to investigate the potential in improvement of genetic gain. Simulated scenarios differed in the method of selection and in the structure of the breeding program. For the base scenario (Sc. Phen-Y12) limited genetic gain could be realized over 25 cycles which in practice would requires 36 years to be completed. The reason for the limited genetic gain is basically due to the fact that (a) this breeding scheme only uses SYN2 groups as parents upon their availability for a new cycle (this means that 11 initial cycles are running independently by using initial varieties as parents and as a result, improvement in genetic gain is delayed and can only be realized by cycle 12 and onward), (b) with phenotypic selection, selection of top *F*_2_ families can be somehow accurate, depending to the *h*^2^ of the trait, however the single plants at this stage are selected randomly from these top *F*_2_ families. Random selection is inefficient in selecting individuals with the highest breeding value as there may be a low genetic correlation between the performance of a plot and single plant being chosen randomly. Thus, the overall genetic gain observed by the end of cycle 25, with this scenario, depends mainly on the accuracy of selection at SYN2 groups (Figure 6).

An alternative for the base scenario (Sc. Phen-Y12) was Sc. Phen. In this scenario, a new cycle could use the *F*_2_ single plants of previous cycles as starting material as soon as they were available. Compared to Sc. Phen-Y12, this scenario realized higher genetic gain by cycle 25 mainly because of shortening the length of breeding cycle. Nevertheless, the realized genetic gain was somehow limited with this scenario due to constraints mentioned above for the selection of *F*_2_ single plants.

In general, genomic breeding schemes had higher genetic gain than corresponding phenotypic based scenarios. The cumulative genetic gain in Sc. GS-Y12 for trait 3 was approximately 2 times greater than the one achieved with Sc. Phen-Y12. Similarly, Sc. GS and GS-SP realized ~3.2 times more genetic gain than Sc. Phen (Figure 4). This indicates that adding genomic information improves selection accuracy of parents that in turn resulted in increased genetic gain. The improvement of genetic gain by GS scenarios is essentially due to more accurate selection for both single plants and plots. In fact, with GS scenarios, the selection of *F*_2_ single plants based on GEBV was efficient and equivalent to having a strong genetic correlation between single plants and plot traits, making it an immensely powerful tool to increase genetic gain. As previously mentioned, random selection of *F*_2_ single plants from *F*_2_ families is not very useful due to lack of genetic correlation between the performance of single plants and their corresponding plot. In addition, the accuracy of selection in SYN2 groups with GS scenarios was more than twice than the accuracy with phenotypic scenarios for initial cycles and accuracy improved over the cycles resulting in more accurate selection of SYN2 groups in later cycles. Thus, basically with GS scenarios not only the genetic gain within each specific cycle was higher than phenotypic scenarios (Figure 5) but also the cumulative genetic gain over the cycles was higher due to improvement in prediction accuracy of both *F*_2_ single plants and SYN2 groups.

Comparing Sc. GS and GS-SP, our results did not show any advantage of using SYN2 single plants (Sc. GS-SP) over SYN2 groups (Sc. GS) as parents for a new cycle. The reason for the similar genetic gain would be that, in both scenarios up to cycle 11, starting material for a new cycle were recruited from the *F*_2_ single plants and theoretically both scenarios would perform the same up to cycle 11. By cycle 12, even though a new cycle could use the output of cycle 1 (either SYN2 single plant or SYN2) as initial material, but apparently *F*_2_ single plants could get higher ranking in terms of GEBV and could be used as initial material for the cycles after cycle 11. As a result, the performance of both scenarios were similar in terms of genetic gain.

Iwata et al. (2011) investigated the performance of GS against phenotypic selection under single trait selection using simulation and found that larger levels of gain could be achieved for the less heritable traits. In contrast, Lin et al. (2016) found larger gains to be achieved for the traits with higher *h*^2^ under multi-trait selection. In our study, we could not realize a dependency between traits *h*^2^ and genetic gain mainly because of the index selection rather than a single trait selection. However, trait 4 with *h*^2^=0.2 had comparable genetic gain as Trait 3 with *h*^2^=0.4 because of a high genetic correlation between these two traits (*r_g_* =0.7).

To evaluate genetic gain, the cumulative genetic gain in all scenarios were standardized by the standard deviation of the *F*_1_ true breeding values at cycle 1 as representative of an unselected stage and cycle. However, the alternative could be to measure genetic gain at the stage of SYN groups as the final product of the breeding cycles. To evaluate genetic gain at SYN stage, then would require standardization by standard deviation of the SYN true breeding values. The problem of standardizing using the SYN variances is that such a variance is influenced by the change in gene frequencies during selection plus the selection during each crossing cycle that led to the SYNs. Therefore, genetic gain was measured at *F*_1_ stage for all scenarios. Alternative measurements of cumulative genetic gain for breeding cycles are in Supplementary Figure 1 and 2.

### Accuracy

The accuracy of selection in SYN2 groups was constant over the cycles with phenotypic scenarios and lower than the accuracy obtained by GS scenarios. Across the cycles, in genomic scenarios the accuracy improved for *F*_2_ single plants, SYN2 groups and SYN2 single plants due to the increase in the size of the reference population. Increasing the reference population size will increase the accuracy achieved (Daetwyler et al., 2010; Albrecht et al., 2011).

Compared to SYN2 single plants, the accuracy was higher in *F*_2_ single plants (Figure 7) despite having smaller reference population at each cycle. As an example, the GEBV for *F*_2_ single plants at cycle 1, were based on the estimated marker effects using the 250 *F*_2_ families of cycle 1, whereas, the GEBV for SYN2 single plants at cycle 1 were based on the estimated marker effects using *F*_2_ families of cycle 1 to 8 and SYN2 groups of cycle 1. The difference in accuracy between two stages can be explained as following. (a) for *F*_2_ single plants essentially training and validation is on the same stage (i.e, the same generation) while SYN2 single plants are two generation away from the training population (random mating of *F*_2_ single plants to create SYN1 and mating within SYN1 groups to create SYN2). In fact, SYN2 single plants are more genetically distant to the reference population compared to *F*_2_ single plants. A general decrease in accuracy in response to an increasing genetic distance between the training and the validation population have been confirmed in livestock genomic evaluations (Habier et al., 2007, 2010). (b) For the initial cycles, the input material for starting a new cycle was from the initial varieties, so even though, the training size for calculation of GEBV in SYN2 single plants, for cycle 1 as an example, was larger but in fact, the extra *F*_2_ families being used for the training, were from independent cycles. In this sense, a large reference population consisting of *F*_2_ families with different genetic makeup cannot translate to a high prediction accuracy in SYN2 single plants.

In our simulation, the reference population was recruited from plot stage *(F_2_* and SYN2) rather than single plants. The use of mean dosage as a genotyping unit per plot is expected to decrease the achievable accuracy in genomic prediction. The reason is that the association of phenotype with mean allele dosage per plot is expected to be less than the association of one phenotype with its individual genotype. The calculation of the mean dosage results in a loss of resolution because the variance of genotypes within the plot population is lost. In this case, large reference populations are required to obtain significant improvements for the accuracy for predicting plot performance. An alternative would be to establish a reference population consisting of genotypes of single plants for the traits that can be reliably measured in single plants (e.g., water soluble carbohydrate content, flowering time). Reference populations would be easier and less costly for collection from spaced plants or clonal rows, especially if phenotypic assessment could be performed with high-throughput methods (Pembleton et al., 2016). It is expected that for such traits prediction accuracy to be higher due to individual assessment of genotyped plants than the traits with reference population consisting of mean dosage as a genotyping unit per plot. Lin et al. (2016) found a higher prediction accuracy (0.7) for flowering time that had a reference population consisting of single plants than productivity traits with a reference population consisted of plots.

In the proposed GS scenarios, the reference population was updated and enlarged in each breeding cycle. This retains the genomic relationship between reference population and selection candidates (i.e., higher probability of SNP being in LD with QTL resulting in better prediction), which is necessary to achieve usable prediction accuracy. Without these updates to the reference population, the accuracy of prediction would deteriorate after a few cycles. This was confirmed by several studies that have investigated the frequency of reference population updates and their effect on genetic gain (Iwata et al., 2011; Yabe et al., 2013).

### Additive genetic variance

Our simulated GS scenarios were found to reduce additive genetic variance over cycles at a more rapid rate than the phenotypic scenarios. Theory predicts a linear decline in additive genetic variance with increasing inbreeding coefficient (*F*) when loci underlying the trait act additively (Buskirk and Willi, 2006). Thus, this reduction of additive variance by GS scenarios indicates that the inbreeding rate per cycle for GS is higher than for the phenotypic selection. Similar results were found by Lin et al., (2017a) where GS resulted in greater inbreeding levels than conventional breeding. These results suggest that control of the extent of inbreeding should be considered in GS-based ryegrass breeding programs. One such method is optimum contribution selection, which aims to maximize genetic gain while restricting the rate of inbreeding per cycle (Meuwissen, 1997). Controls of inbreeding using this method will probably come at a cost of marginally reduced genetic gain in the short term, while delivering higher gain in the long term (Henryon et al., 2015). Therefore, management of inbreeding levels would be prudent when using GS in perennial ryegrass breeding.

A breeding program needs to be affordable to be implemented, in practice. The cost benefit of a new breeding technology such as GS will depend on the value of the extra gain achieved versus the extra cost incurred through genotyping. GS can be introduced in ryegrass breeding programs to replace phenotypic selection in a variety of ways; therefore, implementation of GS in breeding programs will incur several cost components, i.e., genotyping individual plants and plots, which may be best evaluated as cost per unit of genetic gain. Thus, it is important to determine the increments of gain and cost through the whole system in different scenarios to inform the optimal GS option.

## Conclusions

The present study demonstrated the potential for GS to substantially increase genetic gain, as compared with phenotypic selection, when applied under the design constraints of a commercial ryegrass breeding program. This increased genetic gain with GS was mainly due to (a) the ability to select individual plants for plot traits using GEBVs trained from a plot reference population and more effective selection of *F*_2_ and SYN2 groups using GEBVs (b) Reduction of the duration of breeding cycle times, which led to doubling and trebling genetic gain than the traditional selection. However, reduction in additive genetic variance as an indication of inbreeding levels were higher with GS than with phenotypic selection indicating that active methods to simultaneously manage inbreeding and genetic gain will be required.

## Data Availability

The datasets generated for the current study are available from the corresponding author on request.

## Author Contributions

HE developed the scripts, performed the analyses and prepared the manuscript. JJ coordinated the study. HE. DF, and JJ helped to design the scenarios, participated in interpretation of results and revision of the manuscript. BT and LJ contributed in interpreting the results and reviewed the manuscript. All authors read and approved the final manuscript.

---

**Supplementary Figure 1.**
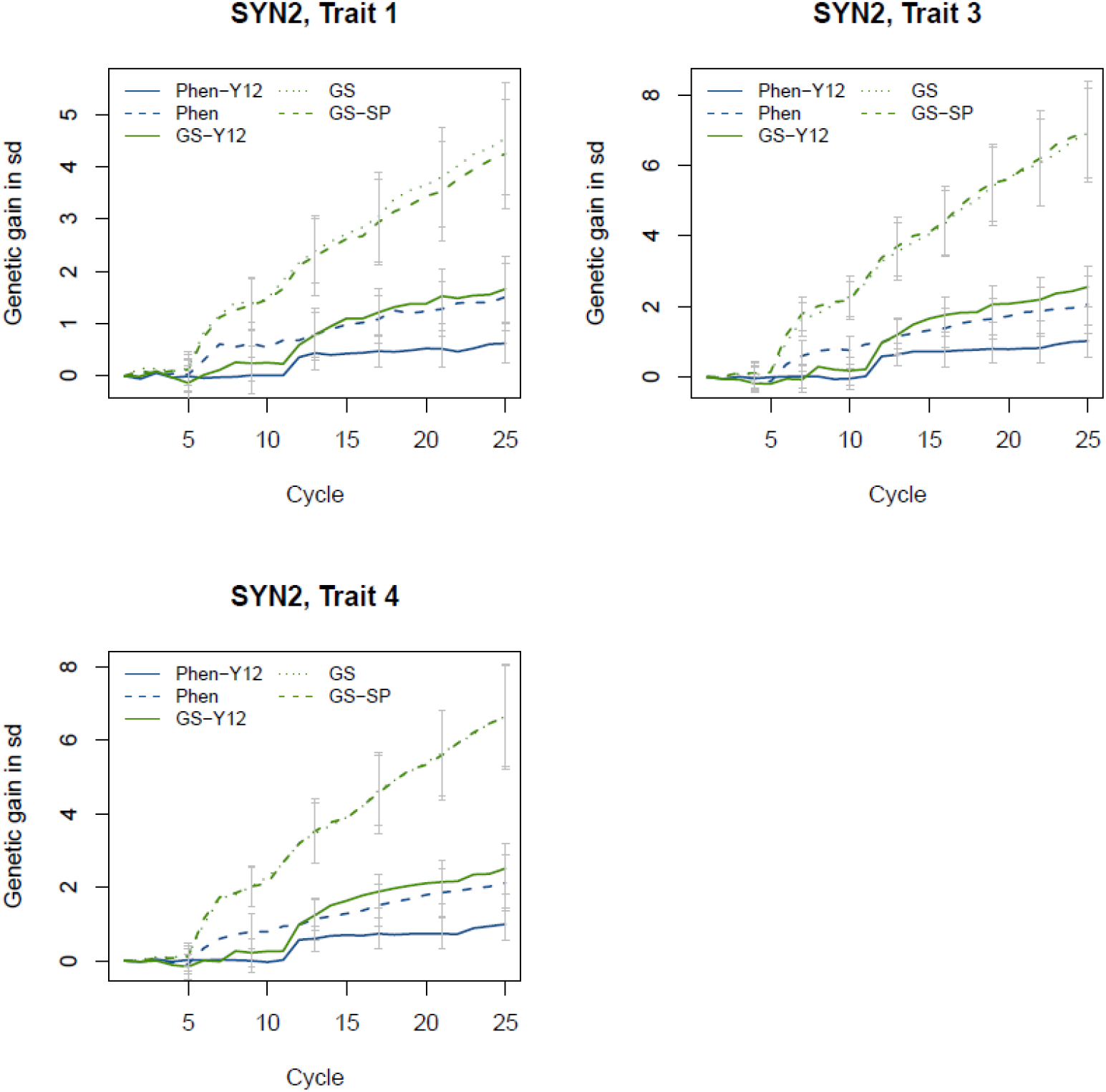
Genetic gain measured at stage of SYN2 and standardized by the standard deviation of the SYN2. Genetic gain was investigated for SYN_2_ stage as:

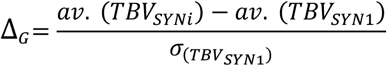

where *av*. (*TBV_SYNi_*) and *av*. (*TBV_SYN1_*) are the mean TBVs of the SYN2 groups in cycle 1 and in later cycles (*i* = 2 to 25), and *σ*_(*TBV*_*SYN*1_)_ is the standard deviation of the SYN2 true breeding values at cycle 1.

**Supplementary Figure 2.**
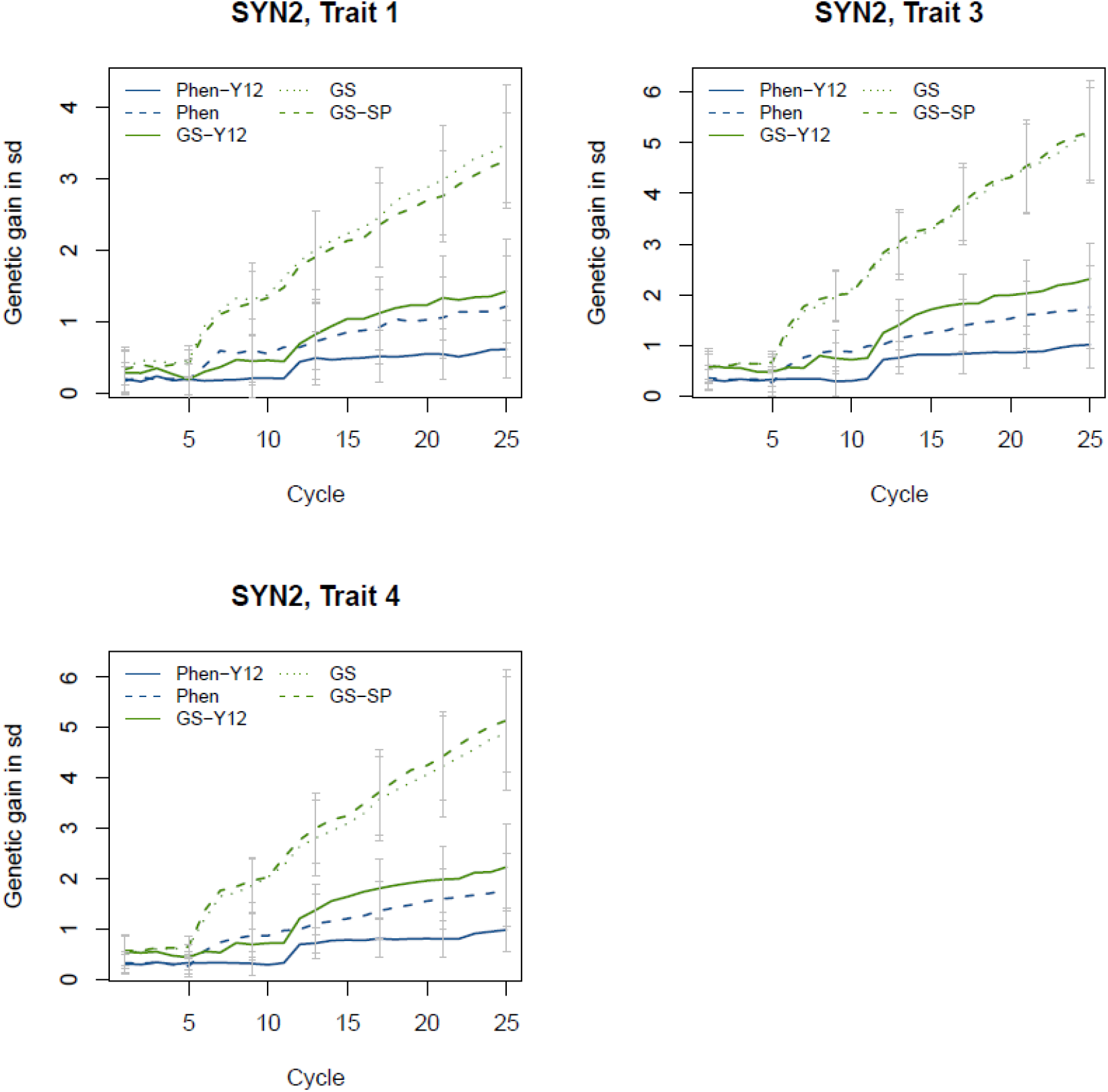
Genetic gain measured at stage of SYN2 and standardized by the standard deviation of the *F*_1_. Genetic gain was investigated for SYN_2_ stage as:

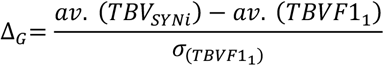

where *av*. (*TBV_SYNi_*) and *av*. (*TBVF*1_1_) are the mean TBVs of the SYN2 and *F*_1_ groups in cycle 1 and in later cycles (*i* = 2 to 25), and *σ*_(*TBVF*1_1_)_ is the standard deviation of the *F*_1_ true breeding values at cycle 1.

